## Supplementary Tables for "First draft genome of the decaploid species, *Ludwigia grandiflora* subsp. *hexapetala*, validated through gene expression"

**Table S1:** Information about genomes used in this study.

| Species | NCBI Accession | Annotation source | Taxonomy (Order/Family) |
|---|---|---|---|
| *Epilobium hirsutum* | GCA_965153245.1 | Annotated in this paper with Helixer | Myrtales/Onagraceae |
| *Chamaenerion angustifolium* | GCA_946814005.1 | Annotated in this paper with Helixer | Myrtales/Onagraceae |
| *Punica granatum* | GCA_007655135.2 | NCBI RefSeq | Myrtales/Lythraceae |
| *Trapa natans* | GCA_035582545.1 | Submitter | Myrtales/Lythraceae |
| *Eucalyptus grandis* | GCA_016545825.1 | NCBI RefSeq | Myrtales/Myrtaceae |
| *Eucalyptus globulus* | GCA_016545825.1 | Submitter | Myrtales/Myrtaceae |
| *Corymbia citriodora* | GCA_016545825.1 | Submitter | Myrtales/Myrtaceae |
| *Rhodamnia argentea* | GCA_016545825.1 | NCBI RefSeq | Myrtales/Myrtaceae |
| *Syzygium oleosum* | GCA_016545825.1 | NCBI RefSeq | Myrtales/Myrtaceae |
| *Syzygium grande* | GCA_016545825.1 | Submitter | Myrtales/Myrtaceae |
| *Melastoma candidum* | GCA_016545825.1 | Submitter | Myrtales/Melasomataceae |
| *Gossypium arboreum* | GCA_025698485.2 | NCBI RefSeq | Malvales/Malvaceae |
| *Citrus sinensis* | GCA_022201045.1 | NCBI RefSeq | Sapindales/Rutaceae |
| *Arabidopsis thaliana* | GCA_000001735.2 | NCBI RefSeq | Brassicales/Brassicaceae |

**Table S2:** Annotation statistics of genes in *Ludwigia grandiflora* subsp. *hexapetala* genome.

| **Annotation type** | **Number of annotation** |
|---|---|
| Total genes | 163,346 |
| rRNA genes | 15,834 |
| 5S rRNA | 2,018 |
| 5.8S rRNA | 3,939 |
| 18S rRNA | 6,237 |
| 28S rRNA | 3,640 |
| tRNA genes | 8,417 |
| Protein coding genes | 139,095 |
| Unique protein coding genes | 122,059 |
| Duplicated protein coding genes | 10,529 |
| Single copy protein coding genes | 111,530 |
| Blast annotations | 103,150 |
| GO annotations | 55,954 |
| Interpro annotations | 79,621 |
| KEGG annotations | 39,714 |
| Uncharacterized proteins* | 53,883 |
| Hypothetical proteins** | 15,767 |

*"Uncharacterized proteins" here correspond to proteins that had a Blast, GO, Interpro or KEGG annotations but without a defined name.
**"Hypothetical proteins" here correspond to proteins that didn't get any of Blast, GO, Interpro or KEGG annotations.

**Table S3:** Predicted subcellular localizations of proteins in *Ludwigia grandiflora* subsp. *hexapetala* genome by WoLF PSORT and LOCALIZER.

| Subcellular localization | WoLF PSORT and LOCALIZER | WoLF PSORT only | Total |
|---|---|---|---|
| Nucleus | 18,924 | 23,085 | 42,009 |
| Chloroplast | 7,957 | 25,019 | 32,976 |
| Cytosol | 0 | 30,022 | 30,022 |
| Mitochondria | 1,990 | 39,92 | 5,982 |
| Plasma membrane | 0 | 5,685 | 5,685 |
| Extracellular | 0 | 5,099 | 5,099 |
| Vacuolar membrane | 0 | 2,193 | 2,193 |
| Endoplasmic reticulum | 0 | 941 | 941 |
| Cytoskeleton | 0 | 830 | 830 |
| Golgi apparatus | 0 | 522 | 522 |
| Peroxysome | 0 | 311 | 311 |

**Table S4:** Repeated elements in *Ludwigia grandiflora* subsp. *hexapetala* genome. Some transposable elements have multiple family annotations due to ambiguous domain, which explains why the total sum of TE is different from the addition of all TE families.

| Classification | Number | Length |
|---|---|---|
| Simple Sequence Repeats (SSR) | 625,820 | 26,674,375 |
| Tandem repeats | 388,071 | 40,880,736 |
| All transposable elements | 140,494 | 57,451,184 |
| Class I/LINE | 9,434 | 4,377,968 |
| Class I/pararetrovirus | 16 | 3,231 |
| Class I/LTR/Ty1-copia | 28,730 | 10,609,375 |
| Class I/LTR/Ty3-gypsy | 98,220 | 40,151,004 |
| Class II/Subclass 1/TIR | 7,070 | 2,904,260 |
| Class II/Subclass 2/Helitron | 797 | 267,502 |

**Table S5:** Number of genes in orthogroups by species. Core refers to genes in orthogroups present in all species.

| Species | Core | Myrtales order only | Family specific | Species specific | Others | Total |
|---|---|---|---|---|---|---|
| *Arabidopsis thaliana* | 25829 | 0 | 0 | 5970 | 13824 | 45623 |
| *Chamaenerion angustifolium* | 21217 | 230 | 366 | 2372 | 12630 | 36815 |
| *Corymbia citriodora* | 24876 | 380 | 623 | 907 | 16317 | 43103 |
| *Citrus sinensis* | 22552 | 0 | 0 | 3006 | 14089 | 39647 |
| *Eucalyptus globulus* | 22394 | 432 | 643 | 499 | 16475 | 40443 |
| *Eucalyptus grandis* | 25104 | 614 | 774 | 853 | 16662 | 44007 |
| *Epilobium hirsutum* | 18404 | 200 | 162 | 266 | 8652 | 27684 |
| *Gossypium arboreum* | 29490 | 62 | 0 | 2652 | 15582 | 47786 |
| *Ludwigia grandiflora subsp. hexapetala* | 50563 | 667 | 464 | 28746 | 44741 | 125181 |
| *Melastoma candidum* | 25858 | 259 | 0 | 3145 | 12312 | 41574 |
| *Punica granatum* | 22078 | 300 | 160 | 1133 | 12237 | 35908 |
| *Rhodamnia argentea* | 25280 | 291 | 437 | 207 | 12325 | 38540 |
| *Syzygium grande* | 16942 | 287 | 340 | 2383 | 12293 | 32245 |
| *Syzygium oleosum* | 24592 | 419 | 536 | 312 | 14350 | 40209 |
| *Trapa natans* | 20670 | 191 | 112 | 404 | 7696 | 29073 |
