## Supplementary Figures for "First draft genome of the decaploid species, *Ludwigia grandiflora* subsp. *hexapetala*, validated through gene expression"

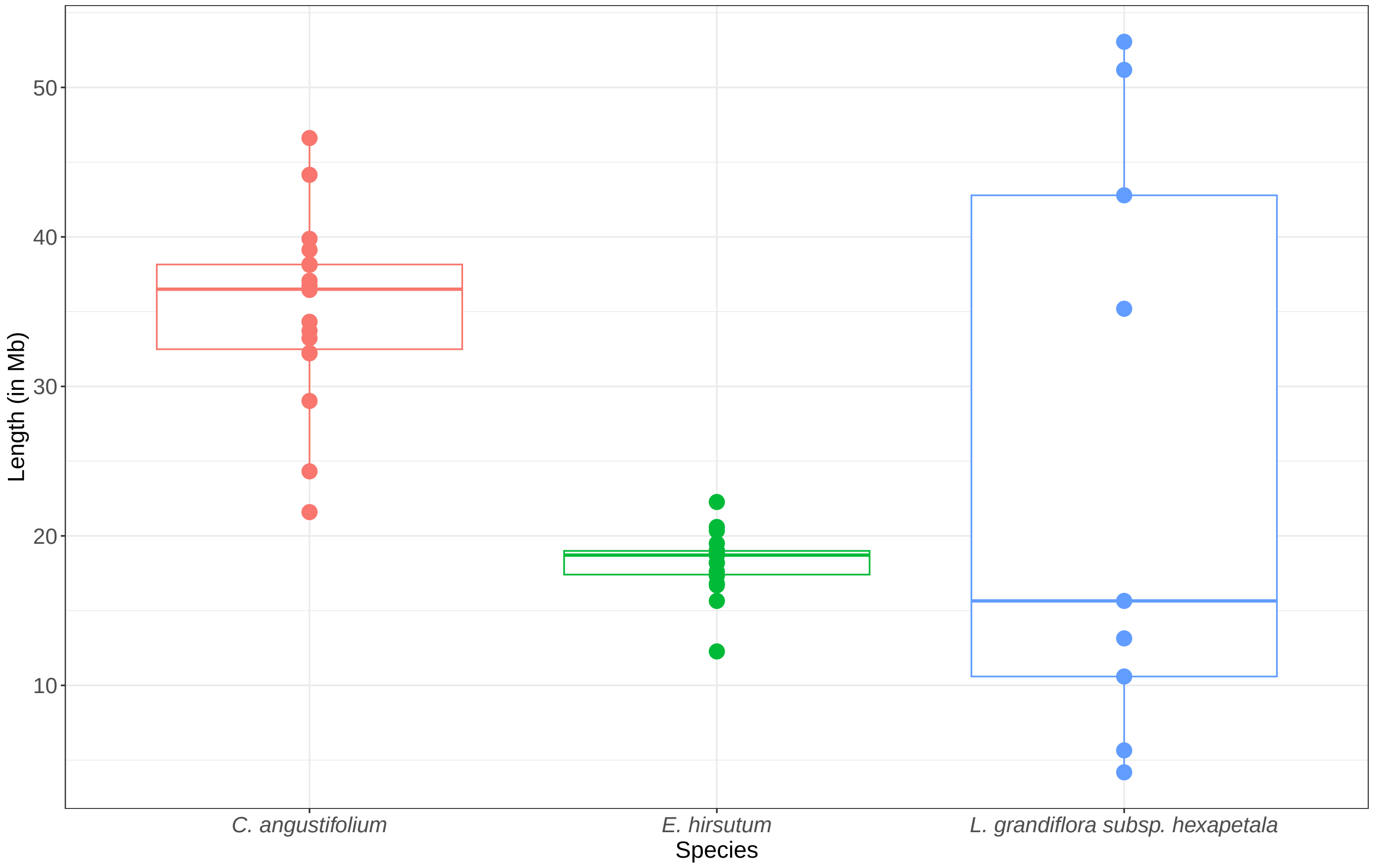


**Figure S1:** Chromosome/scaffold length comparison between *Ludwigia grandiflora* subsp. *hexapetala* (*Lgh*), *Chamaenerion angustifolium* and *Epilobium hirsutum*. Only scaffolds bigger than 4 Mb were represented for *Lgh*.

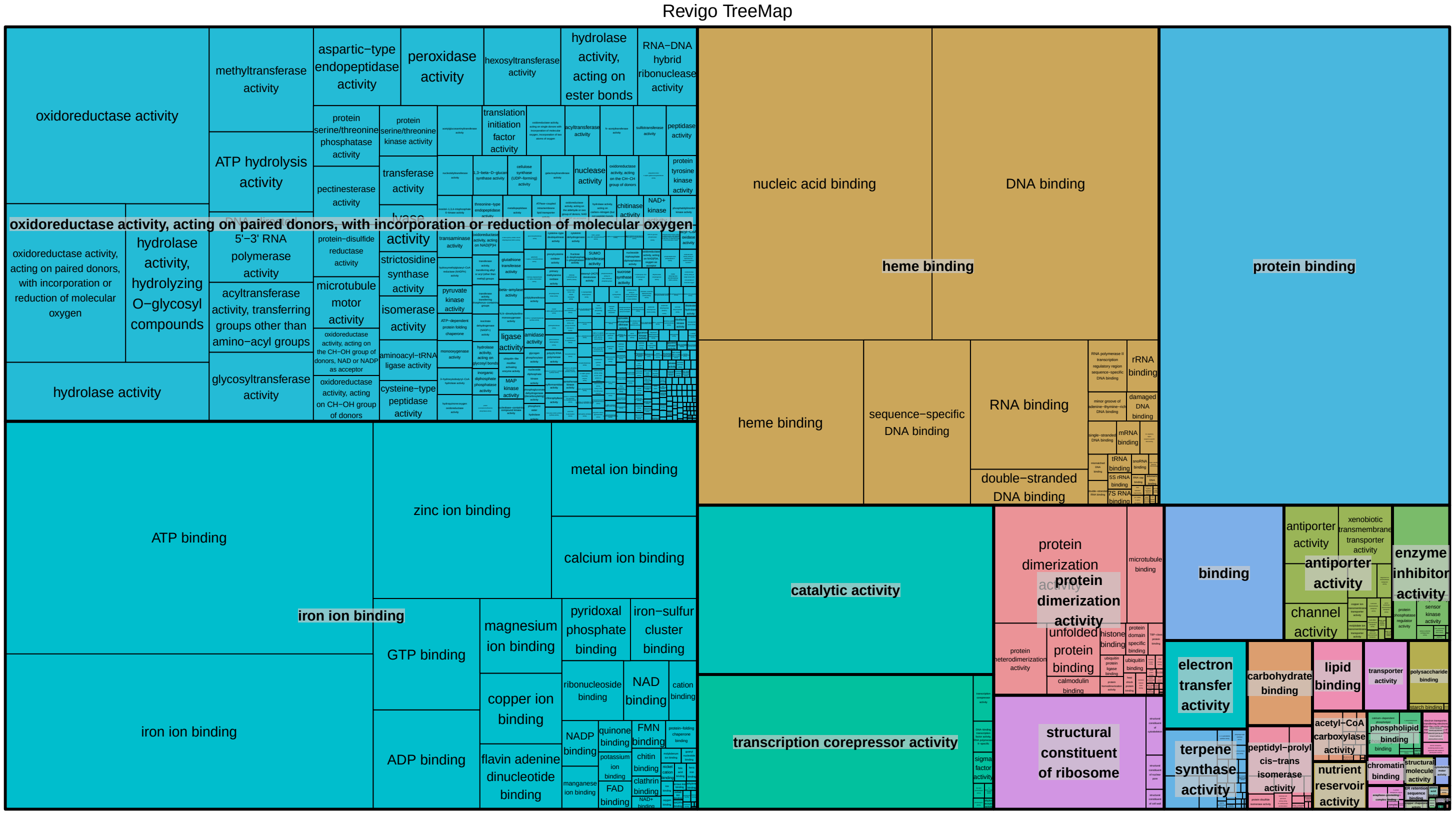


**Figure S2:** REVIGO treemap of GO Molecular function terms based on occurrence frequency in *Ludwigia grandiflora* subsp. *hexapetala*.

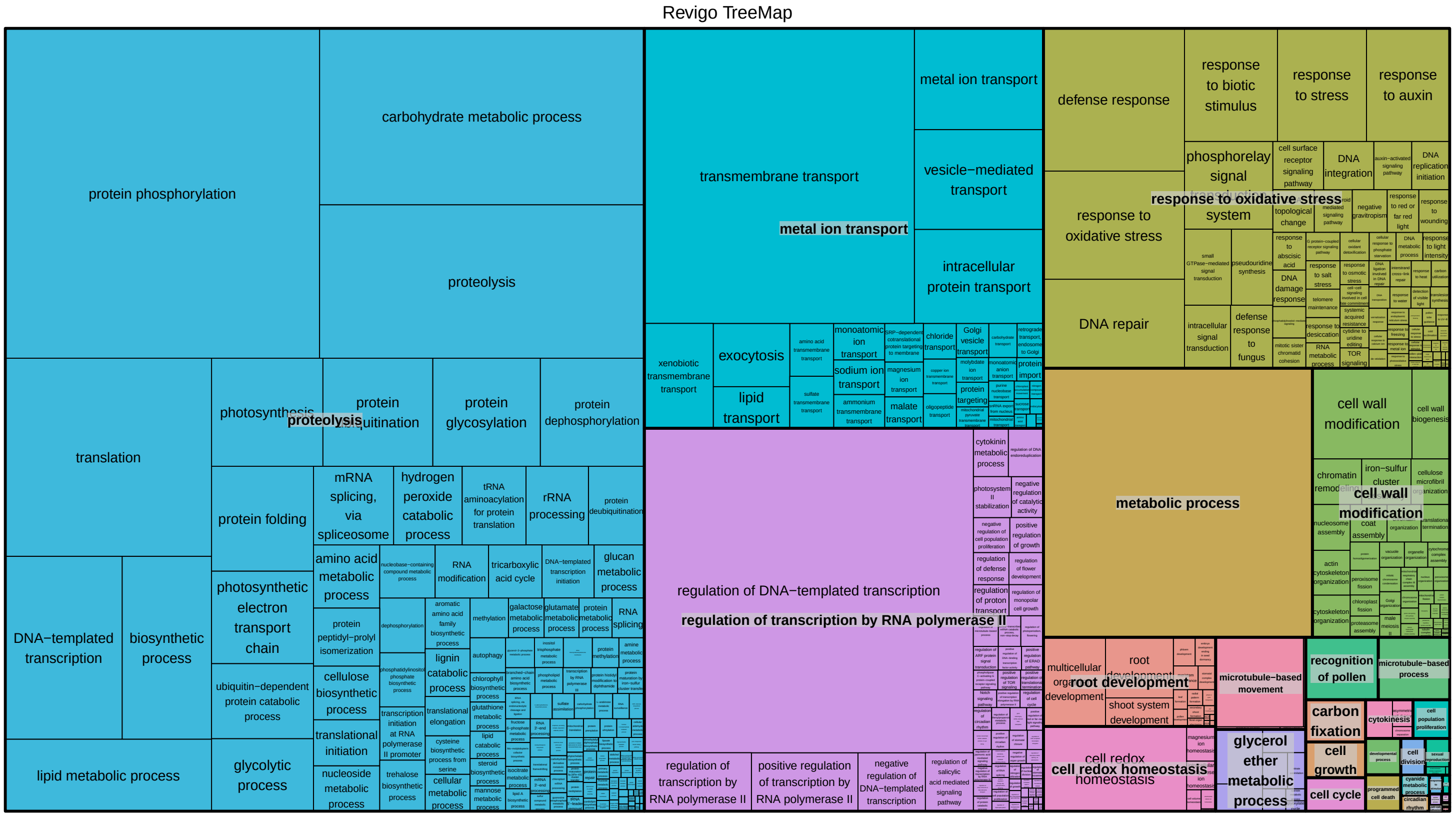


**Figure S3:** REVIGO treemap of GO Biological process terms based on occurrence frequency in *Ludwigia grandiflora* subsp. *hexapetala*.

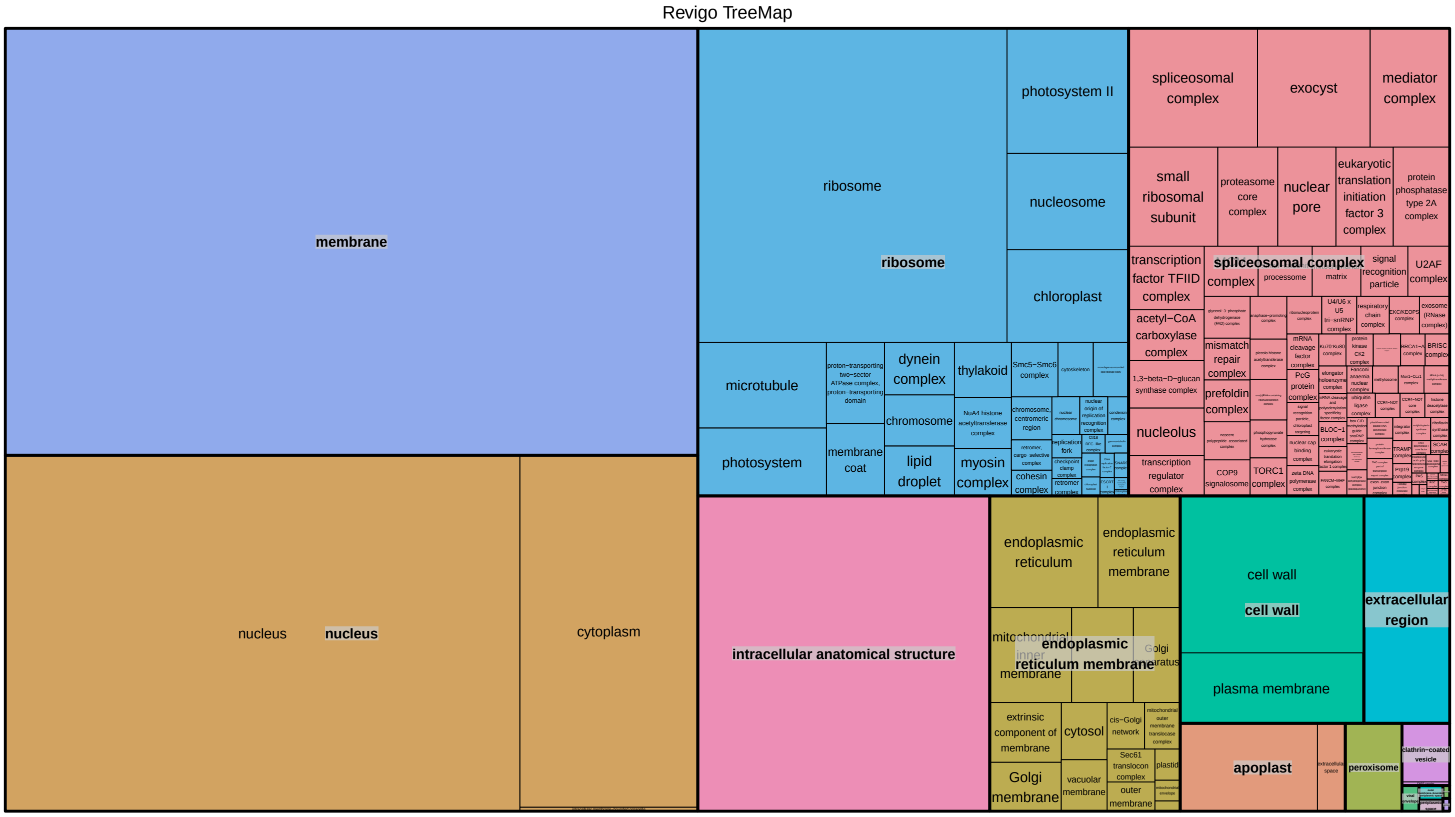


**Figure S4:** REVIGO treemap of GO Cellular component terms based on occurrence frequency in *Ludwigia grandiflora* subsp. *hexapetala*.

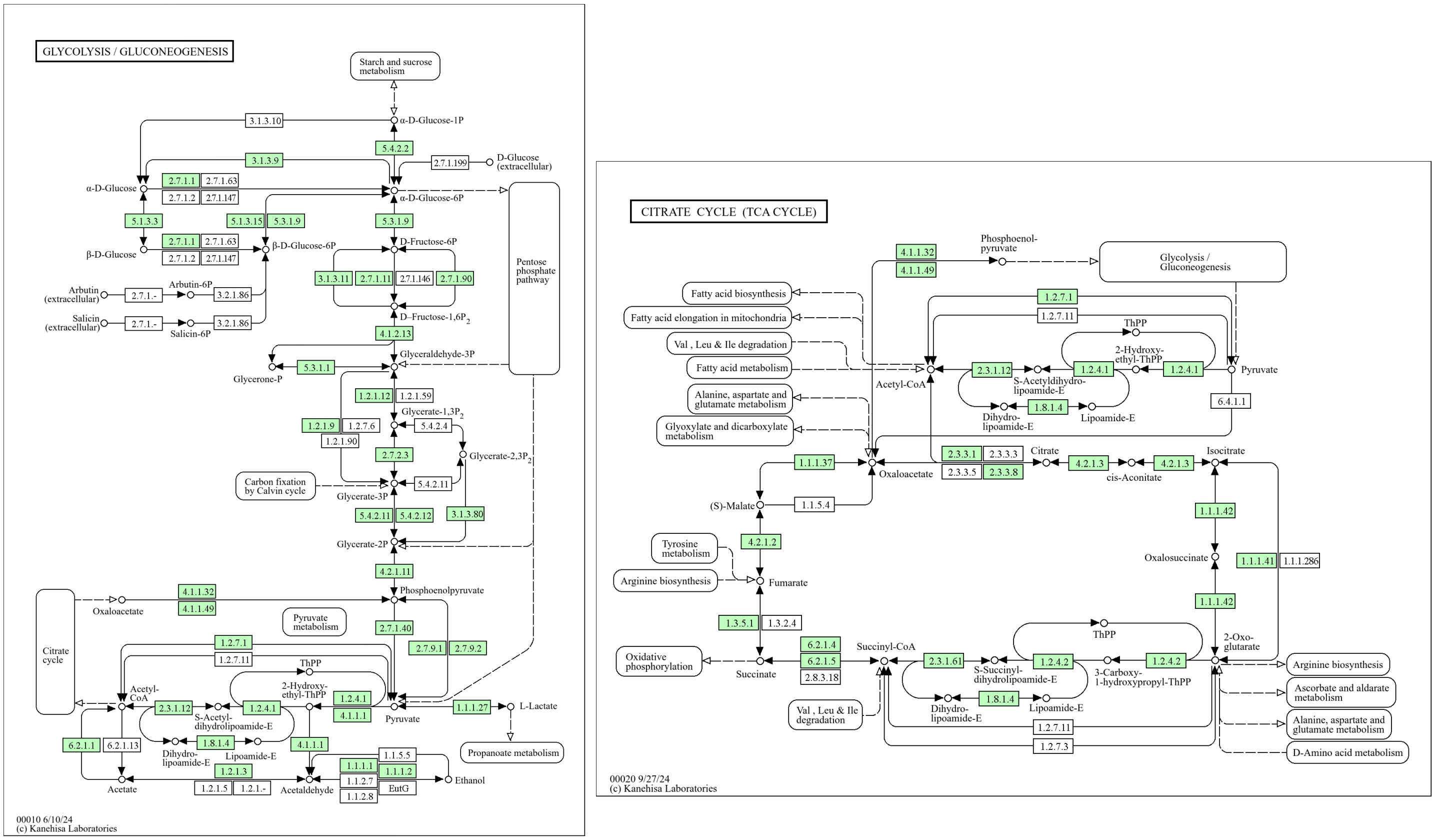


**Figure S5:** Glycolysis and citrate cycle genes in *Ludwigia grandiflora* subsp. *hexapetala* (*Lgh*) genome by KEGG annotations. Genes in green are present in *Lgh* genome.

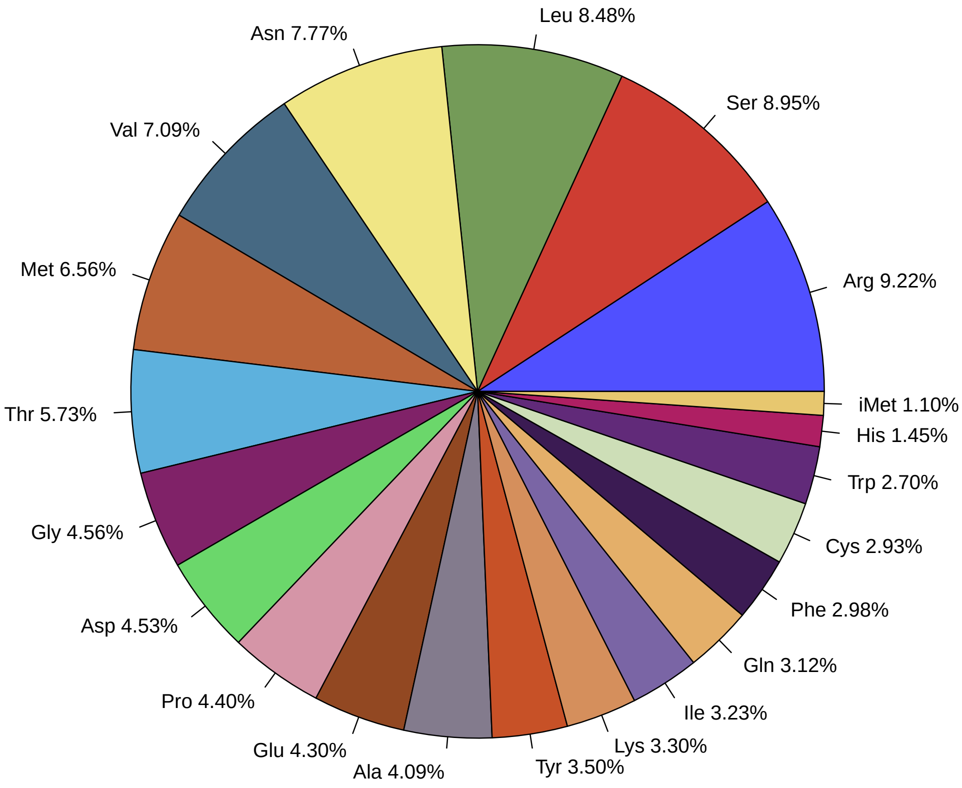


**Figure S6:** Transfer RNA gene repartition by isotype in *Ludwigia grandiflora* subsp. *hexapetala* genome.
